# p57Kip2 regulates embryonic haematopoietic stem cell numbers by controlling the size of the sympathoadrenal progenitor pool

**DOI:** 10.1101/2021.09.13.459835

**Authors:** Chrysa Kapeni, Leslie Nitsche, Alastair M. Kilpatrick, Nicola K. Wilson, Kankan Xia, Bahar Mirshekar-Syahkal, Camille Malouf, Berthold Göttgens, Kristina Kirschner, Simon R. Tomlinson, Katrin Ottersbach

**Affiliations:** Centre for Regenerative Medicine, University of Edinburgh, Edinburgh, EH16 4UU, UK; Department of Haematology, Wellcome Trust-Medical Research Council Cambridge Stem Cell Institute, University of Cambridge, Cambridge CB2 0AW, UK; Institute of Cancer Sciences, University of Glasgow, Switchback Road, Glasgow, G61 1BD, UK; CRUK Beatson Institute for Cancer Research, University of Glasgow, Switchback Road, Glasgow, G61 1BD, UK

**Keywords:** haematopoietic stem cells, aorta-gonads-mesonephros, p57Kip2/Cdkn1c, catecholamines, sympathetic nervous system, neural crest, niche, single-cell RNA-Seq, mesenchyme

## Abstract

Haematopoietic stem cells (HSCs) are of major clinical importance, and finding methods for their *in vitro* generation is a prime research focus. We demonstrate that the cell cycle inhibitor p57Kip2/Cdkn1c limits HSC numbers by restricting the size of the sympathetic nervous system (SNS) and the amount of HSC-supportive catecholamines secreted by these cells, specifically in the aorta-gonads-mesonephros (AGM) region via β2-adrenergic receptor signalling. This regulation occurs at the SNS progenitor level and is in contrast to the cell-intrinsic function of p57Kip2 in maintaining adult HSCs. Using single-cell RNA-Seq we dissect the differentiation pathway of neural crest cells into SNS cells in the AGM and reveal that they are able to take an alternative differentiation pathway, giving rise to a subset of mesenchymal cells expressing HSC-supportive factors. Neural crest cells thus appear to contribute to the AGM HSC niche via two different mechanisms: SNS-mediated catecholamine secretion and HSC-supportive mesenchymal cell production.

## INTRODUCTION

Haematopoietic stem cells (HSCs) with the ability to multilineage repopulate adult recipients are first detected at embryonic day (E) 10.5 in the aorta-gonads-mesonephros (AGM) region of the mouse embryo ^1,2^. They are derived from specialised, haemogenic endothelial cells that can transdifferentiate into blood cells via a process termed endothelial-to-haematopoietic transition (recently reviewed in ^3–6^). While this process is thought to occur in several embryonic tissues harbouring major vasculature, such as the head, yolk sac, placenta, vitelline and umbilical arteries, it is the AGM region where HSCs are first detected in robust numbers and where blood formation from endothelium has been observed by live imaging ^7–10^. This suggests that the AGM has a unique environment particularly suited for promoting HSC formation, which is supported by its remarkable ability to expand HSCs and their precursors in aggregate cultures ^11^.

Relatively little is currently known about the AGM haematopoietic niche and, specifically, the cell types and signals that regulate HSC generation, maintenance and egress ^12^. A large panel of stromal cell lines has been derived from the AGM ^13–15^, which have served as useful tools for the identification of HSC regulators ^16–19^; however, the extent to which these represent the AGM microenvironment *in vivo* is unclear. Cells with characteristics of mesenchymal stromal cells have been detected in the AGM at the time of HSC generation, but their precise location and secretome are still unknown ^20^. Tissues on the ventral side of the dorsal aorta, including the ventral mesenchyme and the gut, are known to provide HSC regulatory signals belonging to the Notch, Hedgehog and Bmp pathways ^17,21–26^. HSC-supportive cytokines such as Scf and Thpo have also been detected in the sub-aortic mesenchyme ^25,27^. In addition, our group has previously reported that cells of the sympathetic nervous system (SNS), which develops in the vicinity of the dorsal aorta at the time of HSC generation, promote HSC production through the secretion of catecholamines, under the control of the haematopoietic transcription factor Gata3 ^28^.

Here, we report that the cell cycle regulator p57Kip2 (Cdkn1c) controls the size of the sympathoadrenal (SA) compartment and thus the amount of catecholamines produced in the AGM, which has a direct influence on HSC numbers. We had previously shown that deletion of p57Kip2 leads to an increase in HSCs at E12.5 ^29^, which is in stark contrast to the essential role of p57Kip2 in adult HSC maintenance and quiescence ^30^ and foetal liver (FL) HSC self-renewal ^31^. We now provide further evidence that, unlike in adult HSC regulation, the effect of p57Kip2 on AGM HSCs is non-cell autonomous and occurs by regulating the proliferation of SA progenitors, hence providing another example of the differences in embryonic vs. adult HSC properties and regulation. We also further define the interaction between the developing haematopoietic and sympathetic nervous system and have generated a detailed characterisation of the emerging SA compartment via single-cell RNA-Seq.

## RESULTS

### p57Kip2 deletion expands HSC numbers in the AGM and the early fetal liver

We had previously reported an increase in repopulation activity of E12 p57Kip2-null AGMs and had suggested that this may be the result of a migration defect in HSCs that causes them to be retained in the AGM for longer before relocating to the FL ^29^. We have now carried out further transplantation experiments, and have also detected a trend towards higher HSC numbers in the E11 p57Kip2-null AGM (**Figure 1A**), which, however, is not as marked as at E12 (**Figure 1B**). Furthermore, we observed significantly expanded HSC numbers in the early FL (**Figure 1C**), which argues against a migration defect and instead suggests that the expansion of the HSC pool in the AGM translates into higher numbers colonising the FL. This initial difference, however, disappears over time as HSC numbers in the FL rapidly increase from E12 (**Figure 1D**). Interestingly, p57Kip2 deletion had no effect on HSC numbers in the E11 and E12 placenta and yolk sac (**Figure 1E-H**), despite high expression in the placenta (**Figure 1I**), demonstrating that p57Kip2 is involved in HSC production specifically in the AGM.

**Figure 1:**
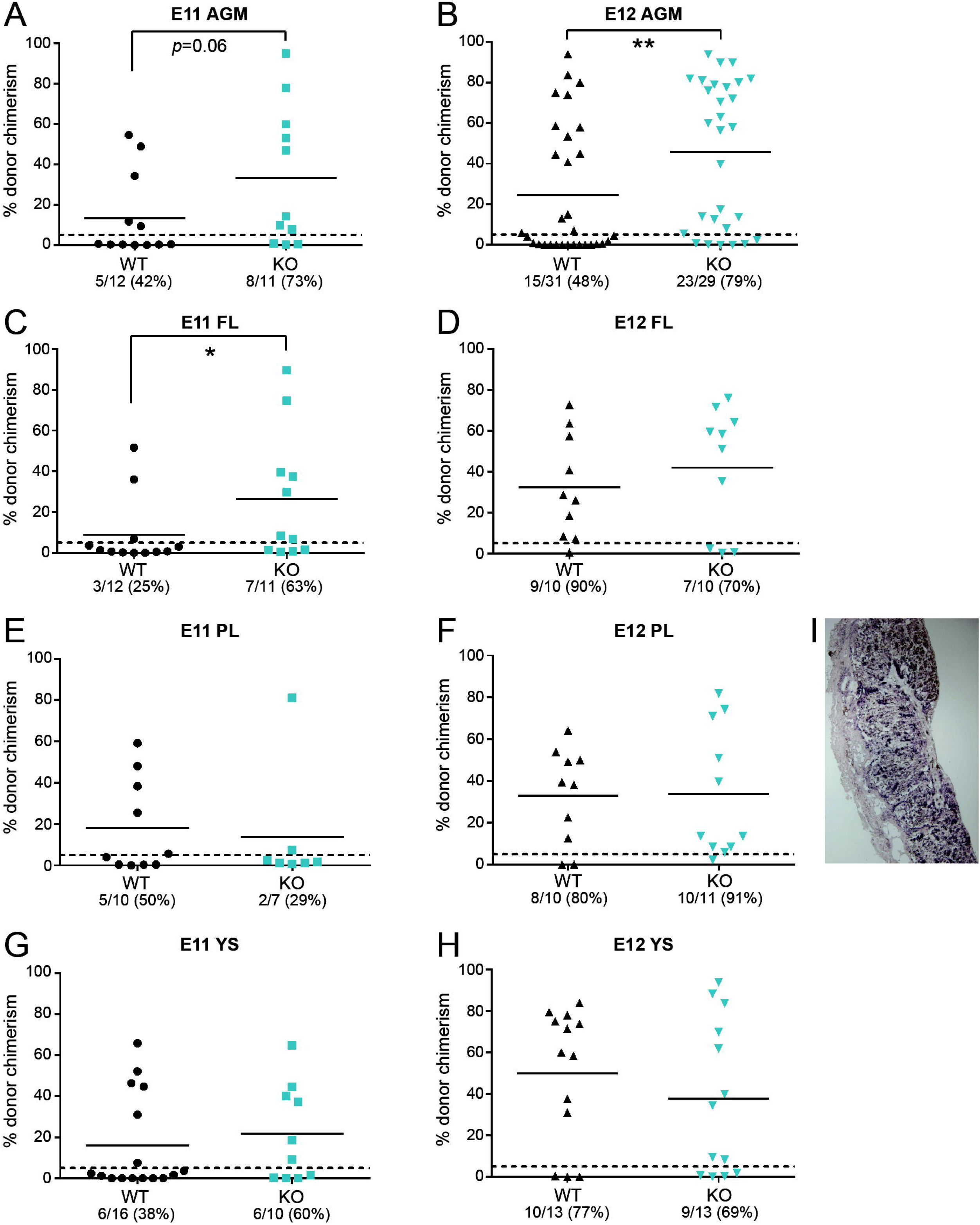
p57Kip2 deletion expands HSC numbers in the AGM and the early fetal liver. Single cell preparations of (**A**) E11 AGM (0.5-1 embryo equivalent [ee]), (**B**) E12 AGM (1ee), (**C**) E11 FL (1ee), (**D**) E12 FL (0.05ee), (**E**) E11 placenta (PL, 2ee), (F) E12 PL (1ee), (**G**) E11 yolk sac (YS, 2ee) and (H) E12 YS (1ee) from wild-type (WT) and p57Kip2-null (KO) embryos were transplanted into irradiated recipients and donor chimerism determined after 4 months by flow cytometry. Data points represent individual recipients with the number of repopulated (>5% chimerism as indicated by the dashed line) recipients/total recipients indicated underneath each graph together with the percentage of repopulated mice. Solid line represents mean; * p<0.05, ** p<0.01, Mann-Whitney *U*-test. (**I**) *In situ hybridisation* with a *p57Kip2* riboprobe on transverse cryosections of E11 placenta showing high expression of *p57Kip2* in trophoblast cells of the placental labyrinth.

### p57Kip2 is highly expressed in the sympathetic nervous system

To understand how p57Kip2 is involved in HSC production in the AGM, we analysed its expression pattern. Immunohistochemistry revealed weak p57Kip2 staining in individual CD34+ endothelial cells (**Figure 2A**, arrowheads) and a slightly stronger signal in subendothelial mesenchymal cells (**Figure 2A,B**, asterisk). By far the most intense staining was observed in patches of cells that were confirmed to be SA cells through co-staining with Ngfr (**Figure 2A,B**, arrows). To obtain more quantitative data, these different p57Kip2+ cell populations were isolated for real-time PCR analysis by fluorescence-activated cell sorting (**Figure 2D, S1**), using CD34 as a marker for endothelial cells, Ngfr for SA cells and Pdgfrβ for mesenchymal cells, excluding SA cells (arrowheads in **Figure 2C**). As p57Kip2 expression had been reported in adult HSCs ^30,32–36^, we also sorted CD45+CD34+ haematopoietic stem and progenitor cells (HSPCs) from the AGM. The expression pattern did not change significantly between E11 and E12 and confirmed the immunohistochemistry data, with the strongest *p57Kip2* expression detected in SA cells and reduced levels in endothelial and mesenchymal cells (**Figure 2E,F**). Considering the reported expression of *p57Kip2* in adult HSCs and its essential role in their maintenance and quiescence, it was surprising to see that there was very little expression in AGM HSPCs. This not only confirms our previous findings that embryonic HSCs have very different properties from adult HSCs, including the way they regulate their cell cycle ^27,28^, but it also suggests that p57Kip2 plays a non-cell autonomous role in HSC production in the AGM.

**Figure 2:**
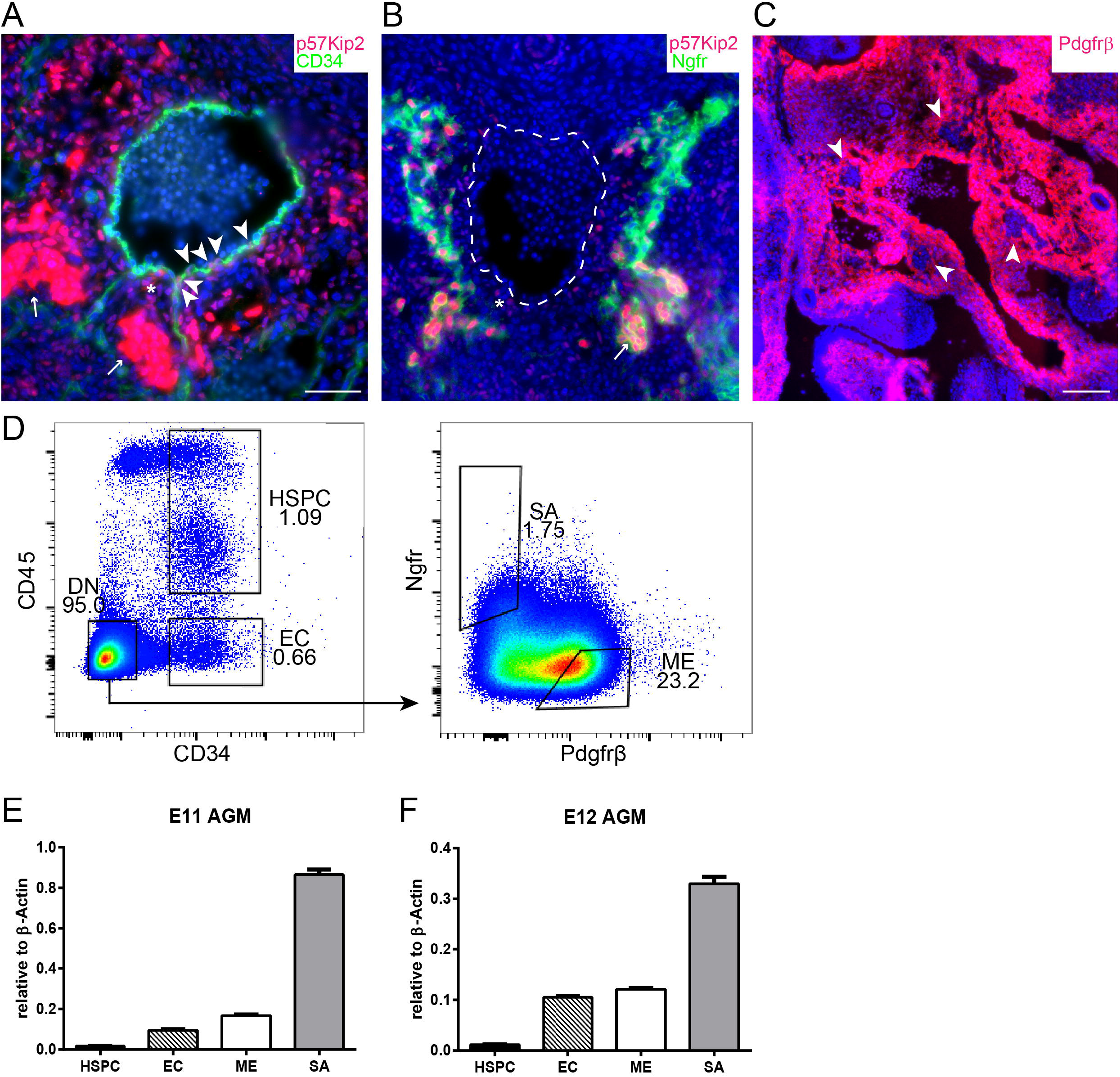
p57Kip2 is highly expressed in the sympathetic nervous system. Immunohistochemistry on cryosections from E11 wild-type embryos. (**A**) Immunostaining for p57Kip2 (red) and CD34 (green) with DAPI nuclear stain (blue). P57Kip2 expression is highlighted in endothelial cells (arrowheads), sub-endothelial mesenchyme (asterisk) and SA cells (arrows). (**B**) Immunostaining for p57Kip2 (red) and Ngfr (green) with DAPI nuclear stain (blue). P57Kip2 expression is highlighted in sub-endothelial mesenchyme (asterisk) and SA cells (arrows). Dashed line shows outline of the aorta. (**C**) Immunostaining for Pdgfrβ (red) with DAPI nuclear stain (blue). Exclusion of Pdgfrβ expression from SA cells is highlighted (arrows). Scale bars indicate 50μm. (**D**) Sorting strategy for AGM subpopulations; HSPCs, CD34+CD45+; endothelial cells (EC), CD34+CD45-; SA cells, Ngfr+Pdgfrβ-; mesenchymal cells (ME), Ngfr-Pdgfrβ+. Gating was based on fluorescence minus one controls (FMOs) as shown in Figure S1. (**E, F**) *p57Kip2* mRNA expression (relative to *β actin*) by qPCR in subpopulations sorted from E11 and E12 AGMs. n ≥ 3. Histogram represents mean ±SEM.

### p57Kip2 increases HSC numbers through an expansion of the catecholamine-secreting SA compartment

Considering the strong expression of p57Kip2 in the SNS and the role of SA cells in promoting HSC production in the AGM via catecholamine secretion, we analysed the effect of p57Kip2 deletion on SNS development. Immunohistochemical staining for the SA marker tyrosine hydroxylase (Th), the enzyme required for catecholamine synthesis, revealed an expansion of the SA compartment (**Figure 3A**), which we had previously observed in *Th* in situ hybridisation experiments ^29^ and have also confirmed by real-time PCR (**Figure S2A**). Interestingly, of all the embryonic tissues harbouring HSCs at that developmental stage, only the AGM expresses robust levels of *Th* (**Figure S2B,C**), suggesting that the functional interaction between the developing haematopoietic and sympathetic nervous system is specific to this tissue. As p57Kip2 is a cell cycle inhibitor, we hypothesised that the expansion of the SA compartment occurs via enhanced proliferation of these cells. Analysis of the cell cycle phases in Ngfr+ cells confirmed a higher percentage of SA cells in the S phase of the cell cycle in the absence of p57Kip2 (**Figure 3B**).

**Figure 3:**
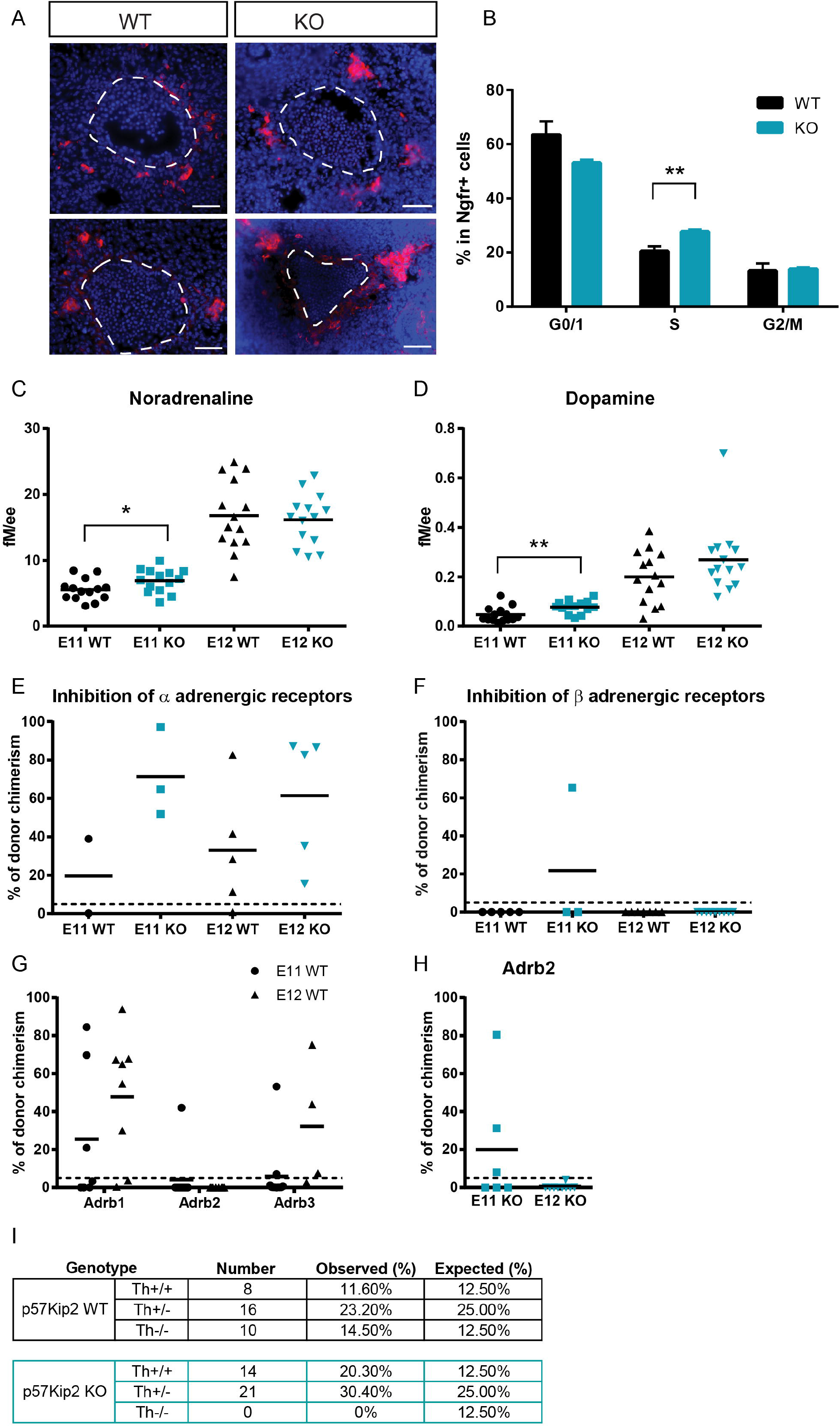
p57Kip2 increases HSC numbers through an expansion of the catecholamine-secreting SA compartment. (**A**)Immunostaining for Th (red) with DAPI nuclear stain (blue) on cryosections from E11 p57Kip2 wild-type (WT) and knockout (KO) embryos. Dashed line shows outline of the aorta. Scale bars indicate 50μm. (B) Percentage of Ngfr+ SA cells from E11 p57Kip2 wild-type (WT) and knockout (KO) embryos in the different cell cycle stages. DAPI-stained Ngfr+ E11 AGM cells were analysed by flow cytometry. (**C, D**) Quantification of the catecholamines noradrenaline (**C**) and dopamine (**D**) by HPLC in individual p57Kip2 wild-type and knockout E11 and E12 AGMs. Concentration is measured in femtomole per embryo equivalent (ee). Black lines denote the mean, n=14. * p<0.05, ** p<0.01, two-tailed, unpaired t-test. (**E, F**) Donor chimerism in recipients of AGM cells from E11 or E12 p57Kip2 wild-type (WT) or knockout (KO) embryos treated *in utero* with the *α* adrenergic receptor (Adra1 and Adra2) blocker phentolamine (**E**) or the β adrenergic receptor blocker propranolol (**F**). (**G**) Donor chimerism in recipients of AGM cells from E11 or E12 p57Kip2 wild-type (WT) embryos treated *in utero* with specific β adrenergic receptor blockers betaxolol (for Adrb1), ICI 118,551 (for Adrb2) and SR 59230A (for Adrb3). (**H**) Donor chimerism in recipients of AGM cells from E11 or E12 p57Kip2 knockout (KO) embryos treated *in utero* with the Adrb2 blocker ICI 118,551. Data points represent chimerism in individual recipients determined by flow cytometry after 4 months, with the dashed line indicating 5% threshold and the solid line the mean. (**I**) Frequency of embryos with the indicated genotypes (E11-12).

To establish whether the expansion of the SNS would translate into a higher production of catecholamines, which might then explain the increase in HSC numbers in the AGM, we measured catecholamine levels by High Performance Liquid Chromatography (HPLC). A pilot experiment revealed that noradrenaline is by far the most abundant catecholamine in the AGM and that its concentration is much higher there than in other haematopoietic tissues (**Figure S2D**). Noradrenaline levels at E11 were significantly higher in the p57Kip2-null AGMs; however, this had normalised at E12 (**Figure 3C**). Even though levels of dopamine, a precursor of noradrenaline, were much lower compared with noradrenaline (**Figure S2D**), we also saw significantly higher levels in E11 p57Kip2 KO AGMs, with a similar trend still visible at E12 (**Figure 3D**).

Catecholamines exert their effects by binding to adrenergic receptors, of which there are two families, the α and the β family. To establish which family is involved in relaying the HSC promoting effect in the AGM, we treated embryos in utero with antagonists specific to these receptor subclasses. Blocking α adrenergic receptor activity had no effect on HSC numbers in the wild-type AGM and also did not abrogate the HSC expansion observed in p57Kip2 KO AGMs (**Figure 3E**). In contrast, blocking β adrenergic receptors had a profound negative impact on HSC activity in both the wild-type and p57Kip2 KO AGMs (**Figure 3F**). As there are three members of the β adrenergic receptor family, we employed more specific antagonists, which revealed that the β2 adrenergic receptor is required for HSC production in the wild-type AGM (**Figure 3G**). This is also the case in p57Kip2-null AGMs, demonstrating that the effect of p57Kip2 on HSCs requires a functional catecholamine - β2 adrenergic receptor signalling axis (**Figure 3H**). To further prove that the role of p57Kip2 in AGM HSC production is indeed upstream of catecholamine secretion from the SNS, we attempted to cross the p57Kip2 knockout line with a tyrosine hydroxylase (Th) knockout line, in which the enzyme required for catecholamine synthesis is deleted ^28^. Surprisingly, the combination of a p57Kip2-null background with a Th-null background resulted in synthetic embryonic lethality, which excluded further analyses of double knockout embryos (**Figure 3I**).

### p57Kip2 is expressed in sympathoadrenal progenitor cells

To better define the cell type in which p57Kip2 is expressed, we performed antibody co-stainings with established markers of the stages of sympathoadrenal differentiation (**Figure 4A** and reviewed in ^37^). The sympathoadrenal lineage derives from Sox10+ neural crest cells, which migrate from the neural tube to the dorsal aorta at around E10. Upon arrival at the aorta, their commitment to the sympathoadrenal fate is initiated by the upregulation of the master regulatory transcription factor Phox2b. Further maturation involves upregulation of Gata3, which is required for the expression of Th, which allows these cells to become fully mature, catecholamine-producing cells of either the adrenal anlage (ventrally) or the sympathetic ganglia (dorsally).

**Figure 4:**
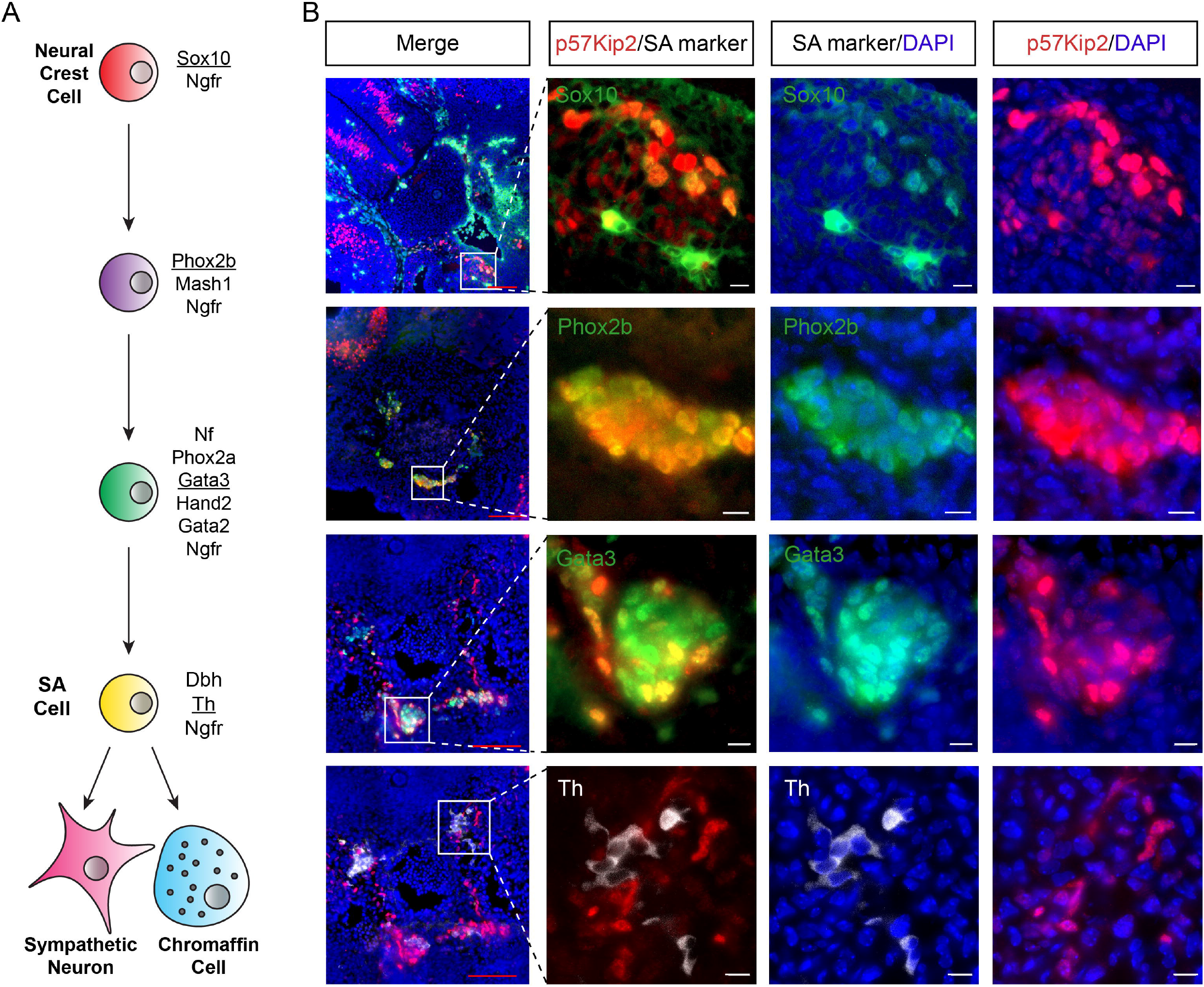
p57Kip2 is expressed in sympathoadrenal progenitor cells. (**A**) Schematic depiction of maturation stages in the SA lineage as defined by listed marker expression. Underlined markers were used in immunohistochemistry in (**B**). (**B**) Immunohistochemical staining of E11 wild-type embryo cryosections for p57Kip2 (red), Sox (green), Phox2b (green), Gata3 (green) and Th (white), with DAPI as nuclear counterstain (blue). Close-ups of boxed areas in merged images are shown. White scale bars equal 10 μm and red bars equal 100 μm.

We detected little overlap between p57Kip2 and Sox10, with p57Kip2 being expressed in cells with a low or no Sox10 signal (**Figure 4B**). Overlap with the master regulator Phox2b, however, is complete. Further down the maturation pathway, p57Kip2 expression becomes more restricted, only partially overlapping with Gata3 and displaying an almost mutually exclusive pattern with Th. These results suggest that p57Kip2 expression initiates with the commitment of neural crest cells to the sympathoadrenal lineage, but that it is gradually downregulated as these cells fully mature to catecholamine-producing cells.

### Single-cell RNA-Seq reveals neural crest differentiation pathways

To better define the differentiation pathway towards a SA fate, we performed single-cell RNA-Seq. As cell surface expression of Ngfr captures all SA cells (**Figure 5A**), we employed this marker to sort single cells from dissociated E11 AGMs by fluorescence-activated cell sorting, excluding mesenchymal cells (Ngfr+ Pdgfrβ-). HSCs are preferentially located on the ventral side of the aorta ^38^. Therefore, to detect factors that are upregulated in the ventral SA domain and that may contribute to the polarised localisation of HSCs, dissected aortae with the surrounding mesenchyme were first cut along the longitudinal axis to separate ventral from dorsal side before single cell preparation and cell sorting. Cell clustering and dimensionality reduction using t-SNE identified six clusters (**Figure 5B**). All clusters contained a mixture of ventrally and dorsally derived cells, apart from cluster 3, which was almost entirely composed of cells with a ventral origin. Cluster 4 appeared to be quite distinct from the remaining clusters and contained cells that displayed virtually no *Ngfr* expression (**Figure 5C**). Instead, these cells expressed high levels of the pan-haematopoietic marker CD45, encoded by the *Ptprc* gene (**Figure 5D**) and other haematopoiesis-associated genes, such as members of the complement system, *Il10ra, Cd52, Cd53* and *Pf4* (**Figure S3A**), hinting at a contamination by haematopoietic cells during the cell sorting. These cells further express high levels of *Csf1r, Cx3cr1* and *Runx1* (**Figure 5E-G, S3A**), but not *Procr* (coding for EPCR), *Kit* or *Cd34* (**Figure 5H-J**), suggesting that they are macrophages resident in the mesenchyme, rather than haematopoietic stem or progenitor cells. Interestingly, *Kit*, normally considered to be a marker of HSPCs, is also expressed in a large portion of SA cells (**Figure 5I**).

**Figure 5:**
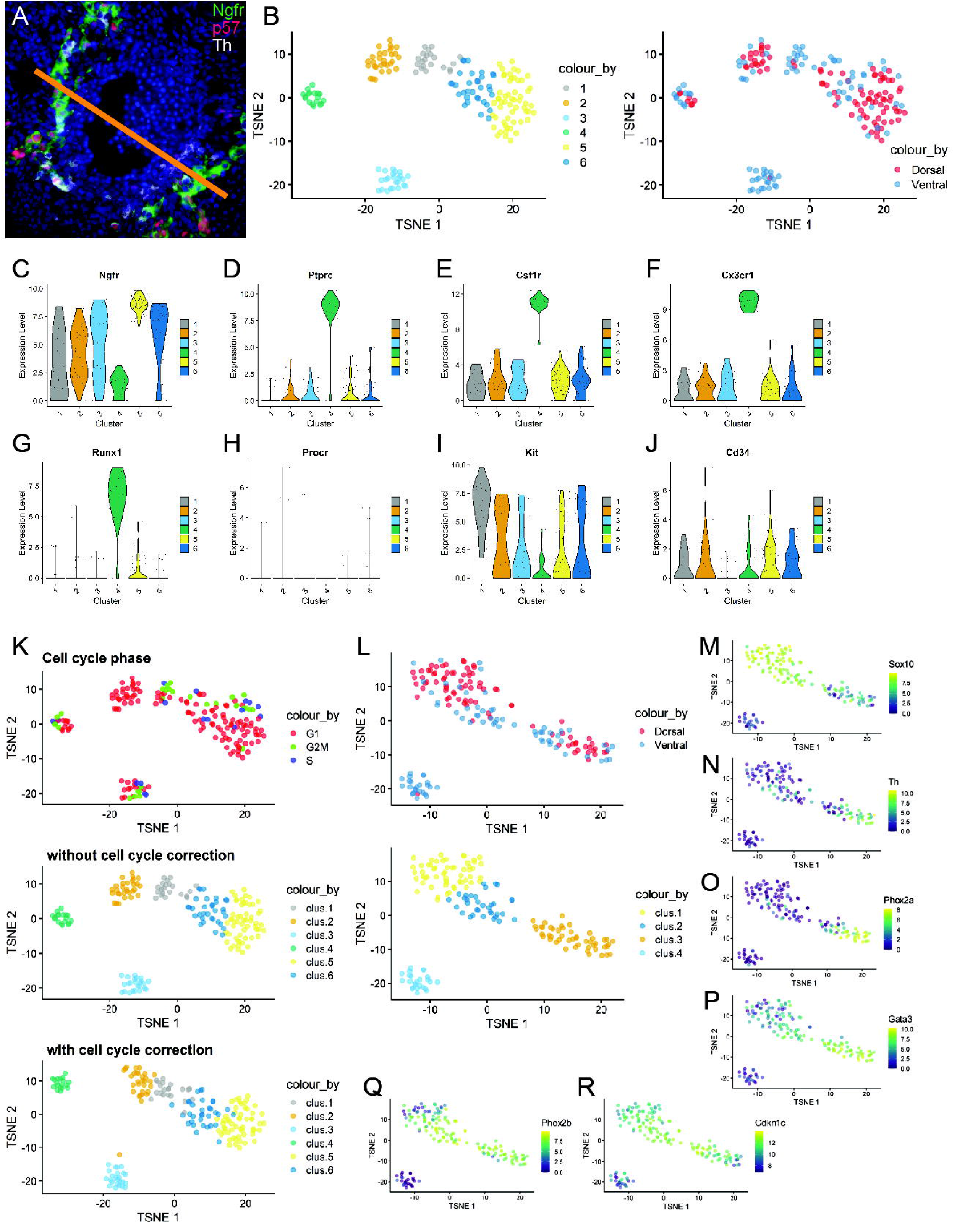
Single-cell RNA-Seq reveals neural crest differentiation pathways. (**A**) Immunohistochemical staining of E11 wild-type embryo cryosection for Ngfr (green), p57Kip2 (red) and Th (white). Bar represents ventral vs dorsal subdissection. (**B**) t-SNE plots coloured for clusters identified by k-means clustering (left) and for ventral and dorsal origin (right). Violin plots for expression levels of *Ngfr* (**C**) and haematopoiesis-associated genes *Ptprc*/*CD45* (**D**), *Csf1r* (**E**), *Cx3cr1* (**F**), *Runx1* (**G**), *Procr*/*EPCR* (**H**), *Kit* (**I**) and *Cd34* (**J**). (**K**) t-SNE coloured for predicted cell cycle phases (top) and the identified clusters before (middle) and after (bottom) cell cycle correction. (**L**) t-SNE plot of remaining 4 clusters following cell cycle correction (bottom) and coloured according to ventral and dorsal origin (top). t-SNE gene expression plots coloured for expression levels of SNS genes *Sox10* (**M**), *Th* (**N**), *Phox2a* (**O**), *Gata3* (**P**), *Phox2b* (**Q**) and *Cdkn1c/p57Kip2* (**R**).

Cell cycle prediction revealed that cluster 2 contained cells entirely in the G1 phase (**Figure 5K**). We therefore applied cell cycle correction, which resulted in the fusion of clusters 1 and 2. After this correction and removal of cluster 4, which contained contaminating macrophages, the remaining cells were now grouped into 4 clusters on which all further analyses were performed (**Figure 5L**). We performed differential expression analysis to determine the 40 most differentially expressed genes for each cluster (**Figure S3B-G**).

To assign the clusters to potential stages of SA differentiation, we investigated the expression of stage-specific markers amongst the clusters (**Figure 4A**). *Sox10* expression was highest in cluster 1 (**Figure 5M**), suggesting that this cluster is enriched for neural crest cells, newly arrived at the dorsal aorta, while *Th* expression, the endpoint of SA differentiation, is mostly confined to cluster 4 (**Figure 5N**). The late SA marker *Phox2a* is also upregulated in cluster 4 (**Figure 5O**), while *Gata3* expression initiates already in cluster 2 (**Figure 5P**). As suggested from the immunohistochemistry staining (**Figure 4**), *Phox2b* and *p57Kip2* show a similar expression pattern. They are both upregulated in some cluster 1 cells, followed by the highest expression in cluster 2 and, at least in the case of *p57Kip2*, downregulation in *Th*-expressing cells of cluster 4. It thus appears that cluster 1 and cluster 4 mark the two endpoints of SA differentiation, with cluster 2 being an intermediate stage.

### Notch signalling is downregulated upon SA maturation

To further confirm this potential differentiation pathway within the developing SA system, we performed cell lineage inference with Slingshot, a method that is specifically designed for modelling developmental trajectories in single-cell transcriptomic data and allows for integration of known developmental stages. Given that the sympathoadrenal lineage derives from Sox10+ neural crest cells, we defined the cluster containing the cells with the highest expression of *Sox10* (**Figure 5M**) as a starting point. Lineage inference analysis revealed that neural crest cells from this cluster (cluster 1) can take two alternative differentiation pathways, ending in clusters 3 (Lineage 2) or 4 (Lineage 1), via an intermediate population (cluster 2), in which divergence of the two fates is initiated (**Figure 6A**). *Ngfr*-expressing cells are found in all clusters, although expression was highest in the neural crest cluster (**Figure 6B**). *Sox10* expression is downregulated along the trajectories, which occurs more gradually towards cluster 4 and more rapidly towards cluster 3 (**Figure 6C**). The trajectory towards cluster 4 appears to be the classic SA differentiation pathway since the end-point was marked by a sharp upregulation of *Th* expression (**Figure 6D**). *Phox2a* is similarly upregulated in cluster 4 (**Figure 6E**), whereas *Gata3* expression, albeit highest in cluster 4, initiates noticeably earlier (**Figure 6F**), which correlates with our immunohistochemistry results (**Figure 4B**). *Gata3* expression is known to be dependent on Phoxb ^39^, the expression of which is clearly initiated prior to Gata3 (**Figure 6G**), again confirming our immunohistochemistry results (**Figure 4B**). Expression of p57Kip2/Cdkn1c is highest in the intermediate cluster 2, going into cluster 4 (**Figure 6H, 5R**), likely making this the stage where SA progenitors are expanded in the *p57Kip2*-null embryos.

**Figure 6:**
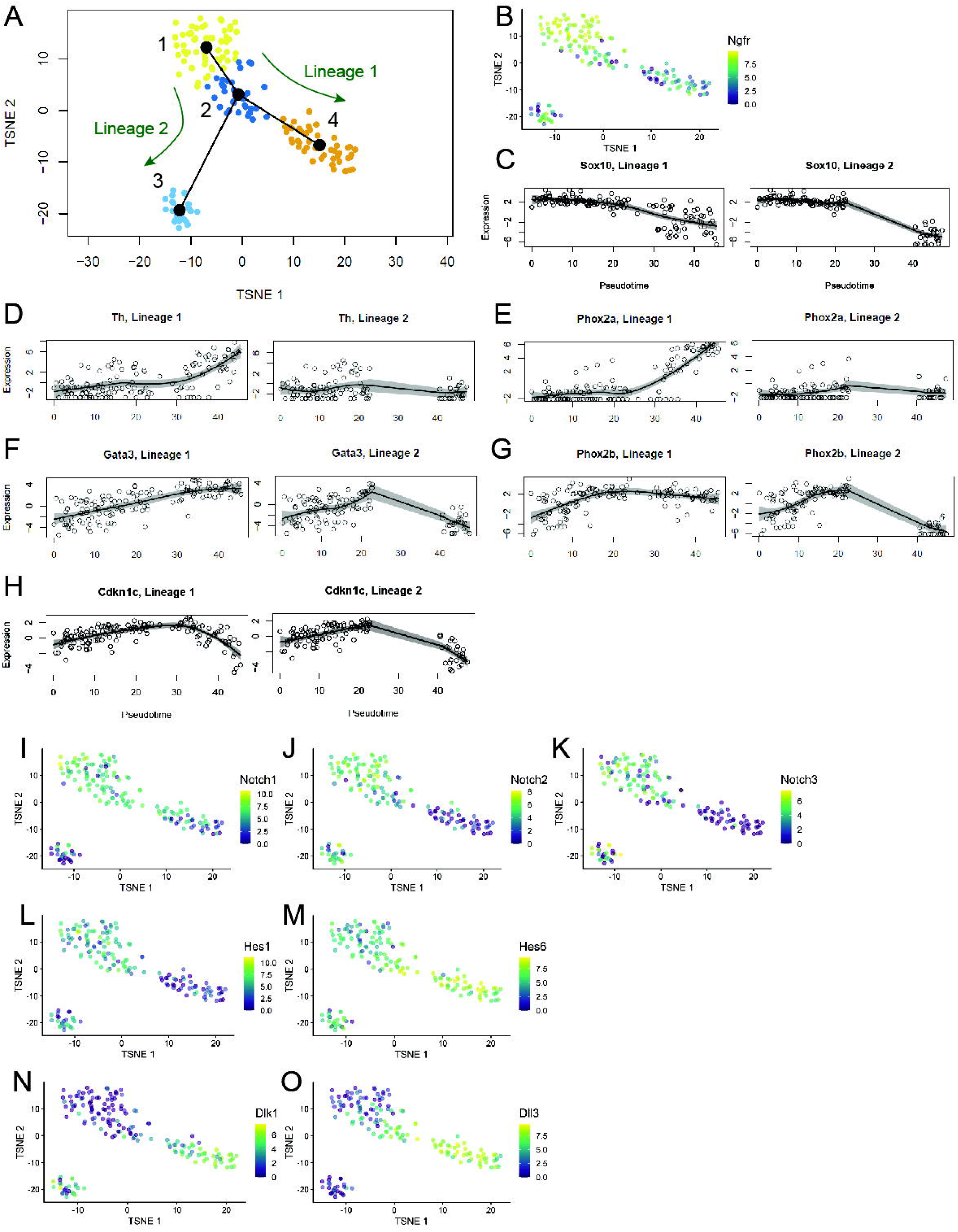
Notch signalling is downregulated upon SA maturation. (**A**) t-SNE plot of 4 clusters with slingshot-identified lineage trajectory nodes superimposed. t-SNE gene expression plot coloured for expression levels of *Ngfr* (**B**). Pseudotime plots for SNS differentiation markers *Sox10* (**C**), *Th* (**D**), *Phox2a* (**E**), *Gata3* (**F**), *Phox2b* (**G**) and *Cdkn1c/p57Kip2* (**H**). t-SNE gene expression plots coloured for expression levels Notch pathway-associated genes *Notch1* (**I**), *Notch2* (**J**), *Notch3* (**K**), *Hes1* (**L**), *Hes6* (**M**), *Dlk1* (**N**) and *Dll3* (**O**).

We noticed that a number of members of the Notch signalling pathway were differentially expressed amongst the clusters (**Figure 6I-O**). The expression of the three receptors, *Notch1-3*, was highest at the neural crest stage, and all three were downregulated by the final stage of SA differentiation in cluster 4 (**Figure 6I-K**). One of the main downstream targets and mediators of Notch activation is Hes1, which showed a similar expression pattern to Notch1-3 (**Figure 6L**). Another member of this family, Hes6, however, displayed the opposite expression pattern, with upregulation coincident with SA differentiation (**Figure 6M**). Interestingly, Hes6 was reported not to be activated by Notch signalling, but instead to serve as an inhibitor of Hes1, thereby promoting neural differentiation ^40^. Two Notch ligands, Dll3 and Dlk1, also showed upregulation along the classic SA differentiation pathway (**Figure 6N,O**), and we have previously shown that *Dlk1* expression in the SNS is Gata3-dependent^24^. Like Hes6, Dll3 and Dlk1 are known antagonists of Notch signalling ^41–44^. Taken together, this suggests that Notch activity maintains neural crest identity and needs to be attenuated for these cells to able to commit to a SA fate, which may be achieved through the upregulation of Notch signalling inhibitors such as Hes6, Dll3 and Dlk1. In support of this, it was demonstrated in chick embryos that expression of the constitutively active intracellular Notch domain inhibited sympathetic neuron differentiation from neural crest progenitors, while inhibition of Notch signalling increased neuron numbers ^45^.

### Neural crest cells can take an alternative differentiation path towards a mesenchymal fate upon arrival at the aorta

Our lineage inference analysis (**Figure 6A**) suggests that, upon arrival at the dorsal aorta, neural crest cells can differentiate down two alternative pathways. As described above, differentiation into cluster 4 cells represents the well-described SA differentiation path as marked by upregulation of key markers such as *Tubb3* and *Dbh* (**Figure S4**). Interestingly, differentiation along the alternative pathway towards cluster 3 cells appears to be spatially restricted to the ventral side of the aorta (**Figure 5L**). The region underneath the ventral endothelium contains a heterogeneous population of mesenchymal cells, many of which secrete factors known to support AGM haematopoiesis (reviewed in ^12^). Furthermore, the existence of cells with properties of mesenchymal stem/stromal cells (MSCs) in the AGM at that time has been reported ^20^. In that context, it is interesting that neuroepithelial cells have been shown to give rise to a transient wave of MSCs via a neural crest intermediate during development ^46^. We therefore hypothesised that a subset of neural crest cells upon arrival at the aorta receives local signals that induce them to differentiate down the mesenchymal lineage instead of the SA lineage. Indeed, a subset of cells in cluster 3 express mesenchymal markers such as Bmper and Pdgfra (**Figure 7A,B**) ^23^. To determine how similar the cluster 3 cells are to sub-aortic mesenchymal cells of the AGM, we integrated a single-cell RNA-Seq dataset from Pdgfrb+ Ngfr-mesenchymal cells isolated from the ventral halves of E11 AGMs. This showed that the putative Ngfr+ mesenchymal cells cluster more closely with the Pdgfrb+ Ngfr-sub-aortic mesenchymal cells, with some individual cells even becoming part of the bigger Pdgfrb+ Ngfr-mesenchymal cluster when projected into a shared t-SNE space (**Figure 7C**). Differential expression analysis of cluster 3 cells compared to all other cells was subjected to Gene Set Enrichment Analysis (GSEA). This further supported the putative mesenchymal character of cluster 3, as significantly enriched Gene Ontology (GO) terms included ‘Heart development’ and ‘Signaling pathways downstream of Tgfb’. To locate cluster 3 cells within the ventral AGM region, we carried out immunohistochemistry analysis of Dlk1, which was upregulated in cells of both cluster 3 and 4 (**Figure 6N**). Co-expression with Th identified cluster 4 cells in ventro- and dorsolateral patches, while Dlk1+ Th-cells were concentrated in the ventral mesenchyme (**Figure 7D**). Bmper has recently been identified as an important regulator for AGM HSC maturation ^23^. We therefore checked for the presence of other important HSC niche factors and detected *Cxcl12* expression in the majority of cluster 3 cells (**Figure 7E**). In situ hybridisation confirmed strong expression of *Cxcl12* in the ventral mesenchyme, while being excluded from the SA domain (outlined in red in **Figure 7F**). In addition, GO terms such as ‘Cytokine-cytokine receptor interaction’ and ‘Chemokines bind chemokine receptors’ were enriched.

**Figure 7:**
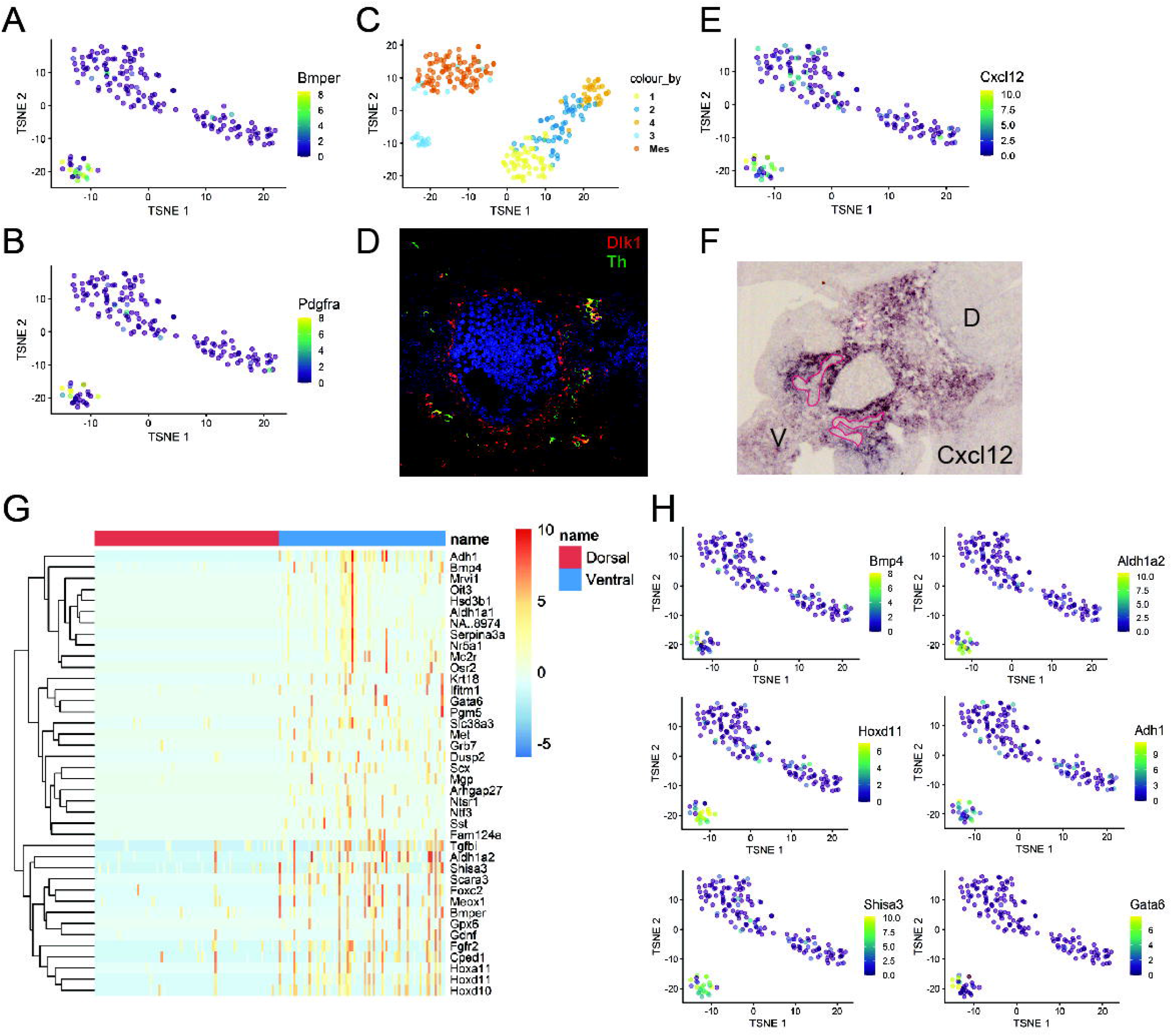
Neural crest cells can take an alternative differentiation path towards a mesenchymal fate upon arrival at the aorta. t-SNE plots coloured for the expression levels of mesenchymal genes *Bmper* (**A**)and *Pdgfra* (**B**). (**C**) t-SNE plot of Ngfr+ cells with Pdgfra+Ngfr-mesenchymal cells sorted from the ventral E11 AGM. (**D**) Immunohistochemical staining of E11 wild-type embryo cryosection for Dlk1 (red), Th (green) and Dapi (blue). (**E**) t-SNE gene expression plot coloured for the expression of *Cxcl12*. (**F**) *In situ* hybridisation staining on E11 wild-type AGM cryosection with a riboprobe for *Cxcl12*. Red lines outline ventral Cxcl12-negative SNS patches; D–dorsal; V-ventral. (**G**) heatmap of the top 40 ventrally expressed genes in Ngfr+ cells. (**H**) t-SNE gene expression plots coloured for the expression of genes upregulated in ventrally derived Ngfr1+ cells.

The detection of HSC regulators prompted us to mine our dataset for other polarised factors, searching for genes significantly differentially expressed between ventrally and dorsally derived Ngfr+ cells. Amongst the genes upregulated in the ventral cells were known HSC regulators, such as *Mgp* ^47^, *Ntf3*^48^ and *Tgfbi*^49^, some of which have previously been shown to be expressed in the ventral mesenchyme, such as *Bmper*^23^ and *Bmp4* ^17,25^ (**Figure 7G**). The expression of the majority of the ventrally upregulated genes was restricted to cluster 3 cells (**Figure 7H**), demonstrating that they are the strongest contributor to the polarised expression of genes within the Ngfr+ populations and may represent an important subset of HSC niche cells. The haematopoietic support function of the ventrally derived cells is further supported by GO terms identified by GSEA, including ‘Peptide ligand-binding receptors’, ‘Factors involved in megakaryocyte development and platelet production’, ‘Systemic lupus erythematosus’ and ‘Retinol metabolism’.

Overall, our results suggest that neural crest cells not only regulate HSC development in the aorta via the secretion of catecholamines from SA cells, which are expanded in the absence of p57Kip2, but that they may also contribute a population of mesenchymal cells to the HSC niche that secrete known HSC supportive factors.

## DISCUSSION

The cell cycle regulator p57Kip2 is strongly upregulated both at the time and in the region of HSC emergence; however, its absence was shown to expand HSC numbers in the AGM ^29^. This study has now further analysed the underlying mechanisms, and we report here that p57Kip2 acts non-cell autonomously through regulation of the size of the SA pool and therefore catecholamine levels. This is in stark contrast to the essential cell-intrinsic role of p57Kip2 in maintaining adult HSC functionality ^30,31^. In fact, we show here that p57Kip2 expression in AGM HSCs is almost absent. This is another example of how embryonic HSCs differ from their adult counterparts in the way they regulate their cell cycle ^28^, respond to DNA damage-inducing agents ^27^ and are affected by mutations linked to haematological malignancies ^27,50^. These differences are clearly important to regenerative medicine and human disease.

This study also delivers further insights into the way in which the definitive haematopoietic system and the sympathetic nervous system interact during their formation in development. Their interaction in the adult bone marrow niche has been well studied, with data supporting a direct effect of catecholamines via the β2 adrenergic receptor on the surface of HSPCs ^51^ or via an indirect mechanism mediated by the expression of the β3 adrenergic receptor on stromal cells ^52^. Our results with adrenergic receptor-specific inhibitors, together with our previous report that the β2 adrenergic receptor is expressed on cells budding from the ventral aortic endothelium ^28^, suggest that the effect of catecholamines (predominantly noradrenaline) is likely to be direct in the AGM. Interestingly, this interplay seems to be restricted to the AGM region, as the p57Kip2 knockout did not affect haematopoiesis in the yolk sac or the placenta and there was comparatively little *Th* expression in these other tissues. This suggests that substantial catecholamine production at this point in development is exclusive to the AGM, as confirmed in our HPLC experiments. The fact that the HSC expansion in the p57Kip2-null AGMs translates into higher numbers of HSCs in the FL at the early stages of colonisation further suggests that the AGM is the main contributor to FL seeding at this stage.

By performing immunohistochemistry and single-cell RNA-Seq, we were able to identify the stage of SA differentiation that is expanded when the p57Kip2-mediated cell cycle break is removed, which corresponds to the Phox2b+ stage following neural crest differentiation, but prior to final maturation as marked by Th expression. Interestingly, expression of p57Kip2 is often associated with terminal differentiation; however, this does not seem to be the case in the SNS, and the absence of p57Kip2 does not appear to impair SNS differentiation. It is therefore puzzling how the combined knockout of p57Kip2 and Th results in such a strong lethal phenotype, especially as either knockout on its own does not result in a decrease in Mendelian ratios before E12.5 (for Th) ^53,54^ and E13.5 (for p57Kip2) ^55^. p57Kip2 knockout embryos are hyperproliferative and may therefore by more acutely dependent on an early supply of catecholamines; however, the exact cause for this synthetic lethality remains to be determined.

The single-cell transcriptome analysis of Ngfr+ cells in the AGM region has provided novel insights into the differentiation pathways that neural crest cells undertake once they have reached the dorsal aorta. In agreement with previous results from chick embryos ^45^, it appears that downregulation of Notch signalling in SA progenitors is required for terminal differentiation. Interestingly, this also seems to be the case for the completion of haematopoietic specification from haemogenic endothelial cells in the AGM ^56–58^, although the mechanisms may differ. Upregulation of Jag was shown to restrict Notch activity in haematopoietic specification, while we observed an upregulation of the Notch signalling inhibitors Dlk1, Dll3 and Hes6.

Intriguingly, our cell lineage inference analysis highlighted a branching point in neural crest cell differentiation with one path leading to SA differentiation (cluster 4), while an alternative pathway results in the generation of cells with a mesenchymal character (cluster 3). There have been several reports of trunk neural crest cells differentiating into mesenchymal cells. An early study suggested that trunk neural crest cells produce the first, yet transient, wave of MSCs ^46^, while subsequent studies provide evidence that at least a subset of bone marrow MSCs, with HSC niche activities, e.g. through the secretion of Cxcl12, are derived from trunk neural crest cells ^59,60^, which may migrate to the bone marrow via the AGM ^61^. Furthermore, a more recent study in zebrafish embryos reported Pdgf signalling-mediated migration of neural crest cells to the dorsal aorta where they promote HSC specification through direct interaction with haemogenic endothelial cells and provision of signals that are catecholamine-independent ^62^. Interestingly, we have also detected upregulation of Pdgf signalling in cluster 3 cells, and they were shown to express HSC regulators such as *Cxcl12* and *Bmp4*. Whether these cells are indeed involved in AGM haematopoiesis and might even interact directly with haemogenic endothelial cells remains to be shown. Cxcl12 and Bmp4 also regulate the migration of neural crest cells and SA progenitors towards the dorsal aorta (reviewed in ^63^), while Cxcl12 and Scf guide primordial germ cell (PGC) migration through the AGM region ^64^. It may thus be an exciting possibility that cluster 3 cells are part of a signalling centre that coordinates the migration/regulation of several important cell types (HSCs, neural crest, PGCs) simultaneously; however, this awaits further studies. Future studies will also aim at identifying the local signals that promote neural crest cell differentiation towards a mesenchymal fate.

## MATERIALS AND METHODS

### Mice and tissue preparations

Animal studies were performed following United Kingdom Home Office regulations. All animals were housed according to institutional guidelines and experiments complied with the animal welfare laws. To obtain embryos, p57Kip2 heterozygous females ^55^ were crossed with C57BL/6J or CD45.1 males. As *p57Kip2* is imprinted, offspring receiving the deleted allele from the mother will have a complete knockout phenotype. For the p57/Th matings, Th+/-males ^28^ were mated with p57+/-Th+/-females. The day of vaginal plug detection was considered E0. Embryos smaller than their littermates or lacking a heartbeat were excluded. Single cell suspensions were obtained by treating AGMs, placentas and yolk sacs with collagenase (Alfa Aesar; 0.125% in phosphate-buffered saline) at 37°C for 45-90 minutes. Foetal livers were dissociated by pipetting.

### Transplantation Experiments

Dissociated cell suspensions of embryonic tissues were intravenously injected into recipients with different CD45 isoforms that had received a split dose of 9.2–9.5 Gy of γ-irradiation. After 1 and 4 months, donor contribution to the recipients’ peripheral blood was determined by flow cytometry, using anti-CD45.1-PE and anti-CD45.2-FITC antibodies (eBioscience). Mice were considered to be repopulated if the donor contribution was at or above 5%. Statistics were performed on GraphPad Prism and the Mann-Whitney *U*-test was used to determine significance levels.

### Adrenergic blockers administration *in vivo*

Each adrenergic receptor was blocked from E8 of gestation. α adrenergic receptors (Adra1 and Adra2) were blocked by intra-peritoneal (i.p.) administration of phentolamine (5 mg/kg). β adrenergic receptors (Adrb1, Adrb2 and Adrb3) were blocked by oral administration of propranolol (0.5 g/l of drinking water) or, for specific β adrenergic receptor inhibition, i.p. administration of betaxolol (1mg/kg) for Adrb1, ICI 118,551 (1mg/kg) for Adrb2 and SR 59230A (5 mg/kg) for Adrb3, all from Sigma. All inhibitors were diluted in PBS, apart from propranolol which was diluted in drinking water and SR 59230A which was diluted in PBS containing 1% DMSO. All blockers were administered every 24h, apart from the β3 adrenergic receptor inhibitor which was administered every 12h.

### Catecholamine detection by HPLC

High Performance Liquid Chromatography (HPLC) was performed in the Psychology Analytical Laboratory (Psychology Department, University of Cambridge). Haematopoietic tissues were dissected and snap-frozen in liquid nitrogen. Perchloric acid (PCA; Fisher Chemicals) was added to the tissues and the samples were homogenised. The samples were spun and the diluted supernatant (1:10 in 0.2 M PCA buffer) was auto-injected into the HPLC machine (234 Autoinjector, Gilson; Hyperclone, 5u, BDS, C18, 100A, 100 × 4.60mm column, Phenomenex). The HPLC system mobile phase consisted of 31.90 g citric acid, 2 g sodium acetate, 460 mg Octanesulfonic acid, 30 mg Ethylenediamine tetra acetic acid (EDTA), 150 ml Ethanol and distilled water up to 1 L; pH was adjusted to 3.6 with NaOH (all from Fisher Chemicals, HPLC grade). The following standards were used to calibrate the HPLC equipment prior to measurement acquisition: NA (noradrenaline), EP (epinephrine/adrenaline) and DA (dopamine), all from Sigma. The signal was detected by an electrochemical detector (ESA, Coulochem II; Parameters: E1 −200 mw, R1 10uA, E2 250 mw, R2 200 nA, flow rate: 0.750 ml per minutes, pressure: 129 bar) and data were collected by the Dionex Data System (Chromeleon). Statistics were performed on GraphPad Prism and unpaired t-test was used to determine significance levels, n=14 per genotype/stage; n=56 in total.

### Flow Cytometry and Cell Sorting

The staining in all the experiments was performed on ice in the dark for 30 minutes. After washing, cells were resuspended in buffer (PBS/2% FCS/ 1% penicillin/streptomycin) containing a viability dye from the Sytox series (Life Technologies) or DAPI (Sigma). All antibodies were purchased from eBioscience, BD and BioLegend, unless otherwise stated. The following antibodies were used: CD45.1 (A20), CD45.2 (104), CD34 (RAM34), Pdgfrβ (APB5), CD45 (104), Ngfr (polyclonal; ANT-007-AO; Alomone Labs Ltd). Cell cycle analysis was performed with DAPI staining. Data were acquired on a Fortessa instrument or cells sorted on Influx or ARIA instruments (all from BD Biosciences). Data were analysed using FlowJo software (vX.0.6, Tree Star, Inc.).

### Immunohistochemistry

Embryos were fixed in 4% PFA (Sigma) in PBS for 1.5 hours at 4°C and equilibrated in 30% sucrose (Fisher Scientific) at 4°C overnight. The following day, the embryos were frozen in OCT (Sakura-Finetek) on dry ice and kept at −80°C until they were sectioned on a cryostat (OTF-5000, Bright Instruments and CM1900, Leica). Blocking solution (2% serum from the animal that the secondary antibody was raised in and 0.4% Triton-X 100 in PBS or 0.1% Tween (all from Sigma)) was added for 1 hour at room temperature (RT). Primary antibodies were then added in blocking solution and left at 4°C overnight. Primary antibodies were: p57Kip2 (rabbit H-91; 1:500, Santa Cruz Biotechnology), Th (mouse LNC1, 1:300, Millipore), Gata3 (goat polyclonal, 1:200, R&D Systems), CD34 (FITC RAM34, 1:200, BD), Phox2b (guinea pig, 1:200, kind gift from J.F. Brunet), Ngfr (goat C-20, 1:300, Santa Cruz Biotechnology) and Sox10 (goat N-20, 1:250, Santa Cruz Biotechnology). The next day, the secondary antibodies were added for 1 −1.5 hours at RT. Secondary antibodies were: Alexa555 goat anti-rabbit (1:500), Alexa647 chicken anti-mouse (1:2000), Alexa488 donkey anti-goat (1:300) and guinea pig (Life Technologies). Subsequently, DAPI solution was added to each slide (1:5000 dilution from 5 mg/μl stocks), which were incubated for 5 minutes. Slides were mounted with 150 μl of Mounting Medium Fluoromount-G (Southern Biotech). The images were acquired on an Axioimager Z2 upright Microscope (Zeiss) (Camera: ORCA Flash 4 v.2, Objectives: 40× Oil and 63x Oil). Extended Depth of Focus images were created from Z-stacks of separate tiles that were then stitched together. Acquisition and processing was performed using the imaging software ZEN 2011 (Zeiss).

### *In situ* hybridisation

Cryosections from frozen embryos prepared as above were air-dried for 20 minutes and fixed in 4% PFA/PBS for ten minutes at room temperature, washed three times with PBS and then acetylated (1.3% v/v triethanolamine, 0.175% v/v hydrochloric acid and 0.25% v/v acetic anhydride in nuclease-free water) for 10 minutes at room temperature. After three washes with PBS, slides were incubated in hybridisation buffer (50% formamide, 5xSSC, 1xDenhardts, 0.02% polyvinylpyrrolidone, 0.02% bovine serum albumin), 0.1% Tween-20, 250μg/ml MRE 600 tRNA, 500μg/ml salmon sperm DNA) for one hour at RT. Digoxigenin-labelled *Cxcl12* riboprobe was diluted 100pg in 100μl hybridization buffer, denatured at 80°C for five minutes and then added to the slides. Slides were covered in parafilm and hybridised overnight at 65°C in a humidified box. The following day, slides were placed in 5xSSC at 65°C to allow the coverslips to detach before four washes in 0.2×SSC at 65°C for 25 minutes each and a final wash for five minutes at room temperature. Slides were then washed in buffer 1 (0.1M Tris at pH 7.5, 0.15M NaCl) for five minutes and pre-blocked in buffer 2 (10% heat-inactivated sheep serum in buffer 1) for at least one hour at room temperature. They were incubated overnight at 4°C with 1:4000 alkaline phosphatase-conjugated anti-digoxigenin antibody (Roche) in buffer 2. On the next day, slides were washed in buffer 1, equilibrated in buffer 3 (0.1M Tris at pH 9.5, 0.1M NaCl, 50mM MgCl2, 0.24mg/ml levamisole) for five minutes at room temperature before being incubated in the dark with 2% Nitro-Blue Tetrazolium Chloride/5-Bromo-4-Chloro-3⍰-lndolyphosphate p-Toluidine Salt (NBT/BCIP; Roche) in buffer 3 at room temperature until staining developed. To stop the staining, slides were rinsed with TE solution (10mM Tris, pH7.5, 1mM EDTA), then water, fixed for 30 minutes in 4% PFA/PBS, then rinsed again with TE followed by water before being mounted in Hydromount mounting medium (National Diagnostics). Images were taken with a Zeiss AxioSkop2 microscope.

*Cxcl12* and *p57Kip2* fragments for riboprobe synthesis were amplified from E11 AGM cDNA by RT-PCR using the following primers: *Cxcl12 fwd*: TTTCACTCTCGGTCCACCTC, *Cxcl12 rev*: TAATTTCGGGTCAATGCACA; *p57Kip2 fwd*: CTGACCTCAGACCCAATTCC, *p57Kip2 rev*: GATGCCCAGCAAGTTCTCTC. The gel-purified fragment was cloned into the p-GEM-T Easy vector (Promega) and digoxigenin-labelled probes generated by *in vitro* transcription using a DIG RNA labelling kit (Roche).

### Gene Expression Analysis by real-time PCR

RNA was extracted using the miRNAeasy Micro kit (Qiagen) according to manufacturer instructions, and RNA quality was assessed by the High Sensitivity RNA Assay (Agilent Technologies) on an Agilent Tapestation instrument. The iScript Advanced cDNA Synthesis Kit for RT-qPCR (Bio-rad) was used according to manufacturer instructions for cDNA synthesis. 2× Sybr (Brilliant III Ultra-Fast SYBR QPCR; Agilent Technologies, UK) was used for qPCR on a LightCycler 480 system (Roche Diagnostics, UK). The program was set as follows: 95°C for 5 minutes, 55 cycles of 95°C for 10 seconds, 60°C for 10 seconds and 72°C for 5 seconds, followed by 95°C for 5 seconds and 65°C for 1 minute. At the end of the qPCR program, a melting curve was run at continuous acquisition mode (97°C, 5 seconds/°C), followed by a cooling step (40°C for 10 seconds). The Ct values were retrieved at the end of the run and data were analysed in Microsoft Excel according to the ΔCt method. All samples were run in at least three biological replicates, each of which was run in technical triplicates. Graphs were made on GraphPad Prism and the student t-test was used to determine significance levels. Primer sequences were: *p57Kip2*: F: 5⍰-CAGCGGACGATGGAAGAACT-3⍰, R: 5⍰-CTCCGGTTCCTGCTACATGAA-3⍰ *Gata3*: F: 5′-CGAAACCGGAAGATGTCTAGC-3′, R: 5′-AGGAACTCTTCGCACACTTGG-3′; *Th*: F: 5′-TATGGAGAGCTCCTGCACTC-3′, R: 5′-TTCTCGAGCTTGTCCTTGGC-3′; *β-Actin*: F: 5′-GGCTGTATTCCCCTCCATCG-3′, R: 5′-CCAGTTGGTAACAATGCCATGT-3′.

### Library preparation for scRNA-Seq

scRNA-seq analysis was performed using the Smart-seq2 protocol as described previously ^65^. Single Ngfr+Pdgfrb- or Ngfr-Pdgfrb+ cells were sorted by FACS directly into individual wells of a 96-well plate containing lysis buffer (0.2% RNase inhibitor (Ambion, Thermo Fisher Scientific) in Triton X-100 (Sigma)), and libraries were prepared using the Illumina Nextera XT DNA preparation kit. Pooled libraries were run on the Illumina Hi-Seq 2500 and reads aligned using STAR ^66^.

### Read alignment

Reads were aligned using Kallisto (Linux v0.43.0, ^67^) using parameters `kallisto quant --plaintext --bias --single -fragment-length=200 --fragment-length=200 -sd=20`.

Samples were mapped to Mus_musculus.GRCm38.cdna.all.fa version of the transcriptome downloaded from Ensembl (www.ensembl.org) Feb. 2017. This library was appended with the ERCC92 spikes (ERCC92.fa) downloaded from www.thermofisher.com and appended to the transcript library. Read quality from individual libraries was assessed using FastQC (https://www.bioinformatics.babraham.ac.uk/projects/fastqc) and MultiQC, ^68^).

Transcripts were trimmed using Cutadapt^69^) using the command `cutadapt -a CTGTCTCTTATA -f fastq -e 0.1 -O 3 -q 20 -m 20`).

### Data pre-processing

Data were pre-processed following the single-cell transcriptomics workflow of Lun et al. ^70^ using R version 4.0.0 under 64bit Windows >= 8. Data were imported into R using the scater Bioconductor package (scater_1.18.6, ^71^) function readKallistoResults. Transcript level quantitation was also collapsed to gene at this stage using a transcript to gene mapping table generated from Ensembl Transcript Stable ID and Gene Stable ID. All downstream analysis was carried out at gene level. Ensembl IDs were mapped to MGI Gene Symbols using Bioconductor mouse gene annotation files (org.Mm.eg.db _3.6.0).

Following the Lun et al. ^70^ workflow, cell quality was assessed based on library size, number of expressed features in each library, percentage of reads aligned to mitochondrial genes (obtained from Ensembl) and percentage of reads aligned to spike-in transcripts. We used the median absolute deviation (MAD) definition of outliers to remove putative low-quality cells from subsequent analysis: cells with any quality measurement more extreme than 3 MADs from the median were removed. Similarly, cells which had outlying proportions of mitochondrial genes or samples with low numbers of unique genes were also removed. Removing cells with a high proportion of reads mapping to spike-in RNAs left only one cell in the HSC1 sample group after filtering. This cell was removed from the data and the HSC1 group eliminated from further analysis. The HSC2 group was referred to as HSC in the rest of the analysis. Overall, 335 cells consisting of 65 HSC2, 93 Dorsal, 89 Ventral and 88 Mesenchymal cells passed initial filtering. We filtered genes with low abundance (those with mean count across all cells less than 1), since these genes are likely to be dominated by drop-out events, limiting their usefulness in further analysis. 14,109 genes passed this filtering step.

We used the computeSumFactors function in the scran R package ^70^ to normalise within each plate and then scale plate samples to each other. Read counts mapping to ERCC spikes were normalised separately to the expression data and not used for normalisation of expression samples. In this analysis we cannot completely exclude confounding plate-effects. However, internal control marker and cell profiles were used to assess the normalisation and are consistent with the expected biology.

### Cell cycle correction

Following the workflow of Lun et al. ^70^, we used the Cyclone prediction method ^72^ to assign cells into cell cycle phases based on the gene expression data. Cells were assigned to a cell cycle phase if the score for that phase was greater than 0.5. Cells were considered of ambiguous cell cycle phase otherwise. Cells were initially visualised using their cell cycle phase before removal of this effect using limma (v3.46.0, ^73^) by fitting a linear model (^~^G1+G2M +S) and decomposing the variance. The trendVar function was used to fit a loess smoothed line to the gene specific variances and the decomposeVar function was used to decompose the variances into biological and technical variances. Genes were selected with the biological component of the variance >=0.5 and FDR <0.2 against the null hypothesis that biological variance equals 0. This generated a set of highly variable genes that was used as a set of genes for dimensional reduction with t-SNE.

### Cell clustering and dimensional reduction

t-distributed Stochastic Neighbour Embedding (t-SNE; ^74^) was used for dimensionality reduction, t-SNE visualisations were generated with varying settings of the perplexity parameter (5, 10 and 20) and the clusters obtained found to be robust. A t-SNE perplexity value of 10 was chosen as representative. t-SNE was performed across all the data and coloured by group: HSC2, Dorsal, Mesenchymal and Ventral cells. A set of predetermined marker genes for Dorsal/Ventral (Phox2b, Ngfr, Sox10, Cdkn1c, Th, Gata3), HSC (Ptprc, Kit, Procr, Runx1, Cd34) and Mesenchymal (Pdgfra, Col1a2, Dlk1, Cspg4, Acta2, Pdgfrb and Cxcl12) were used to confirm the biological identify of each cluster.

A group of cells that were from Dorsal/Ventral origin were selected from the data and used to identify highly variable genes, dimensionally reduced using t-SNE and split into six clusters using k-means clustering. Robustness of the computed clusters was confirmed using the Dynamic Tree Cut method. Analysis of the computed clusters revealed that the cells in one cluster expressed a range of known macrophage markers; cells in this cluster were removed from downstream analysis. Further analysis of the remaining clusters identified one cluster that was purely defined by cell cycle identity; following batch correction for cell cycle using limma and reclustering, cells in this cluster were found to be redistributed among the remaining four clusters.

### Differential expression analysis

The edgeR Bioconductor package (v3.32.1; ^75^) was used to identify significantly differentially expressed genes between cell clusters. Differential expression results were filtered by fold change and an FDR threshold of 0.2. The top 40 differentially expressed genes for each cluster were computed using fold change ranking and a fixed FDR threshold of 0.2.

### Cell lineage inference analysis

The Slingshot Bioconductor package (v.1.8.0; ^76^) was used to perform cell lineage inference analysis. The expression data and cell cluster labels were used as input to infer the global lineage structure. Based on this structure, two smooth cell lineages were constructed and pseudotime variables were inferred for both smooth lineages.

### Gene set enrichment analysis

The GSEAPreranked tool (GSEA 4.1.0, build 27; ^77^) was used for gene set enrichment analysis. To create the ranked gene list, genes were filtered to remove genes with logCPM <= 0; remaining genes were ranked based on log fold change. Gene sets were obtained from the ConsensusPathDB-mouse interaction database ^78^; gene sets containing less than 15 genes or more than 500 genes were excluded from further analysis. The default GSEA enrichment statistic (weighted) was used, corresponding to a value of p=1 in the GSEA enrichment score calculation ^77^. GSEA’s meandiv method (default) was used to normalise enrichment scores to account for differences in gene set size and allow analysis over gene sets. Multiple hypothesis testing was corrected using sample permutation (n=10000).

### Data accessibility

The single cell RNA-seq data from this study have been deposited in the Gene Expression Omnibus database (GEO,^79^) under the accession number GSE139052.

## ACKNOWLEDGMENTS

We are indebted to the staff of the animal facilities both at the Cambridge Institute for Medical Research and the Centre for Regenerative Medicine for their support with animal experiments and to the flow cytometry teams at both of these institutes, Dr. Reiner Schulte, Dr. Chiara Cossetti, and Michal Maj in Cambridge and Fiona Rossi and Dr Claire Cryer in Edinburgh, for excellent cell sorting services and help with flow cytometry analyses. We are also very grateful to Dr Brunet for providing us with the Phox2b antibody. Core facilities at the Edinburgh Centre for Regenerative Medicine were supported by centre grant MR/K017047/1. This work was funded by a Bloodwise Bennett Senior Fellowship (10015 to K.O.) and the Kay Kendall Leukaemia Fund (to K.O.). This research was also funded in part by the Wellcome Trust and the UKRI Medical Research Council. For the purpose of open access, the author has applied a CC BY public copyright licence to any Author Accepted Manuscript version arising from this submission.

## AUTHOR CONTRIBUTIONS

C.K. performed and designed the majority of experiments and wrote the manuscript; L.N., A.M.K. and K.K. performed bioinformatics analyses; N.K.W. and B.G. provided advice and assistance with scRNA-Seq experiment; K.X., B.M.-S. and C.M. performed experiments; S.T. designed, performed and supervised the bioinformatics analysis of the scRNA-Seq data; K.O. conceived and supervised the study and wrote the manuscript.

## SUPPLEMENTAL FIGURE LEGENDS

**Figure S1**

Sorting gates and fluorescence-minus one (FMO) controls for isolating endothelial cells (EC), haematopoietic stem and progenitor cells (HSPC), sympathoadrenal cells (SA) and mesenchymal cells (ME) from the AGM.

**Figure S2**

(**A**) Real-time PCR analysis of *Gata3* and *Th* expression in E11 wild-type (WT) and p57Kip2 knockout (KO) AGMs. *Th* expression by real-time PCR in the AGM, foetal liver (FL), yolk sac (YS) and placenta (Pla) from E11 (**B**) and E12 (**C**)wild-type embryos. (**D**) Analysis of catecholamine levels by HPLC in the adult bone marrow (BM), AGM, FL, YS and Pla.

**Figure S3**

Heatmaps showing the top 40 differentially expressed genes in the macrophage cluster 3 before cell cycle adjustment (**A**) and, after cell cycle adjustments, in the remaining four clusters (**B-G**).

**Figure S4**

Heatmaps showing the top 40 differentially expressed genes in the two alternative differentiation pathways of neural crest cells.

