## Supplementary figures and images for "p57Kip2 regulates embryonic haematopoietic stem cell numbers by controlling the size of the sympathoadrenal progenitor pool"

### Supplemental Figures

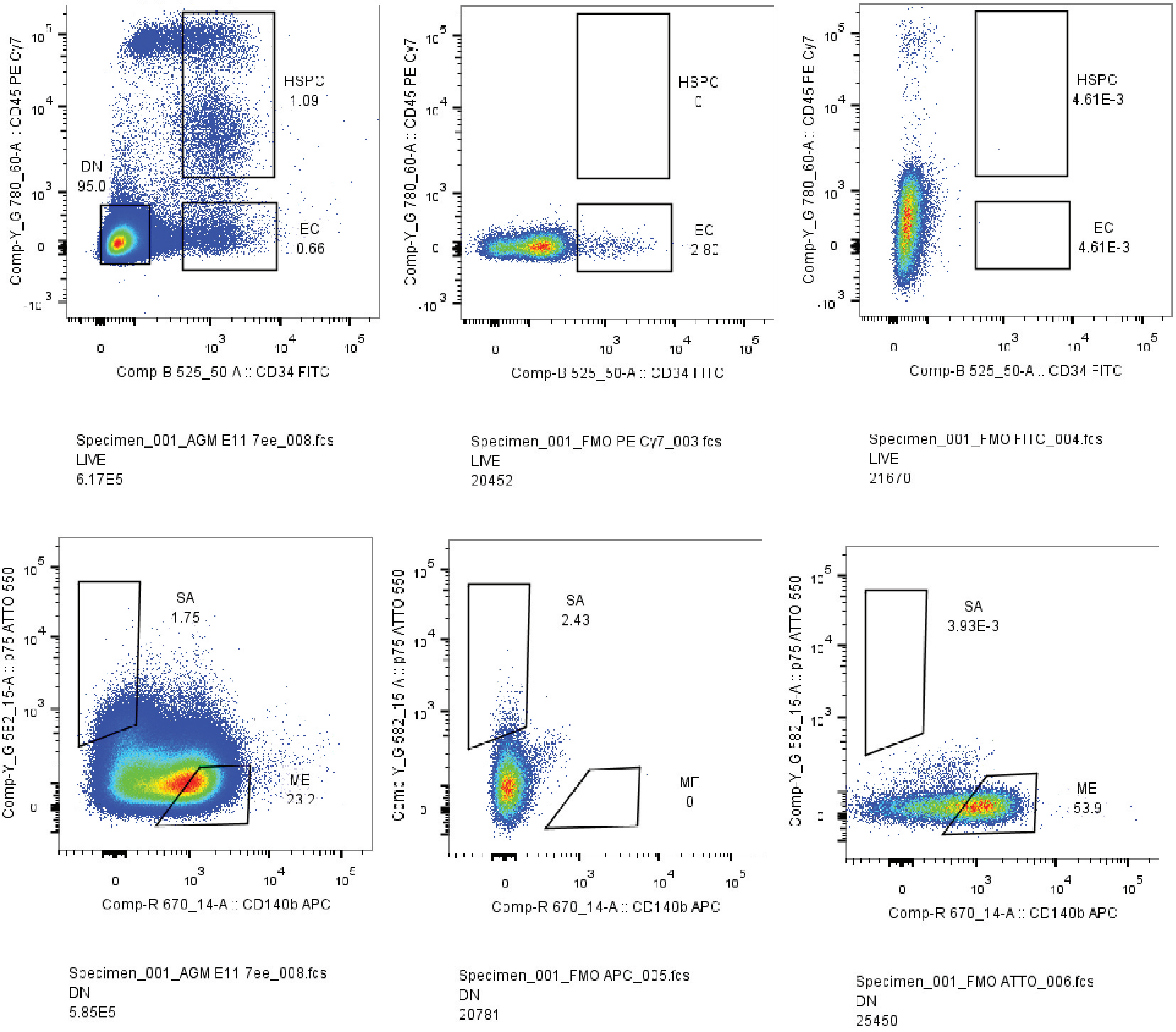

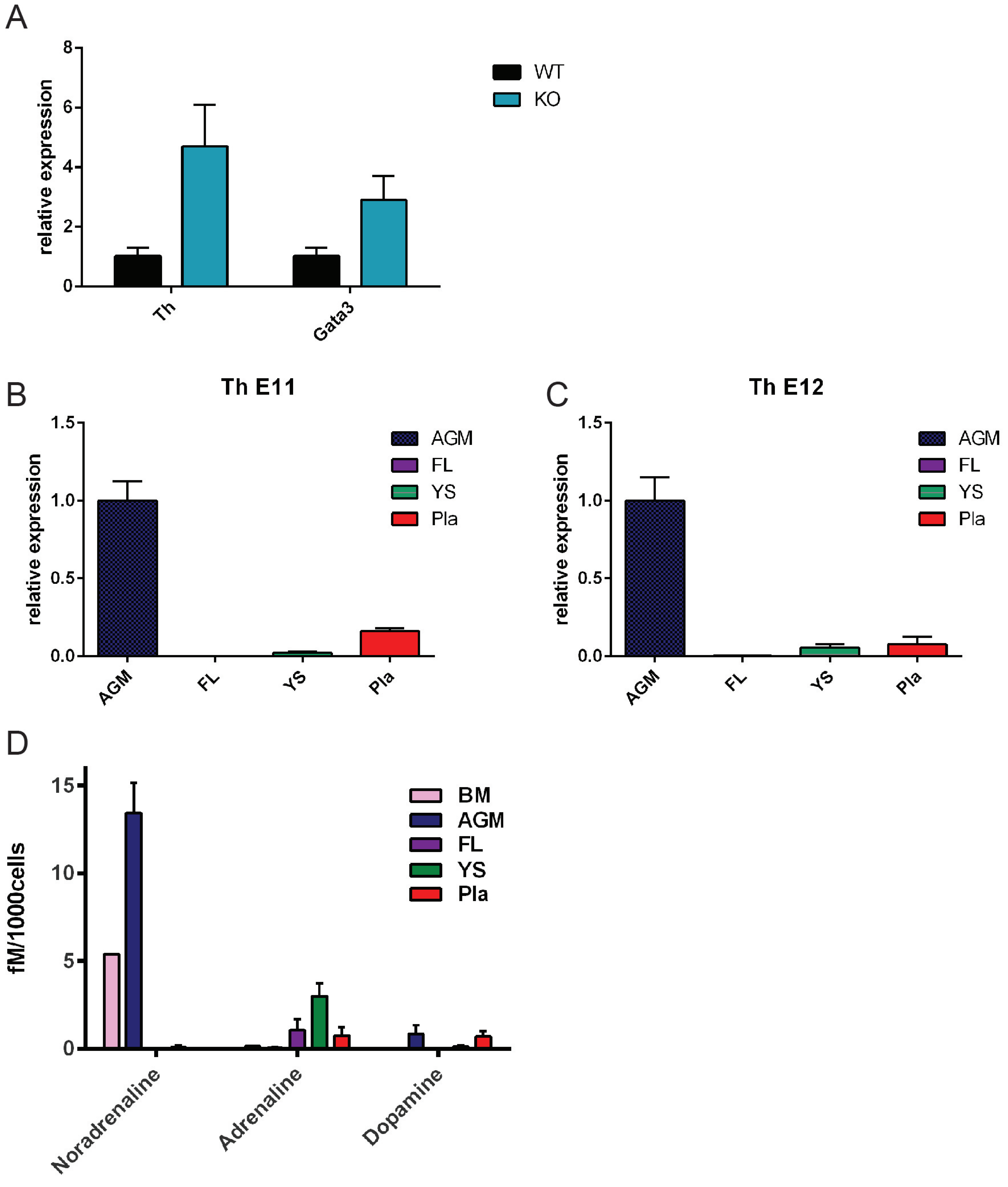

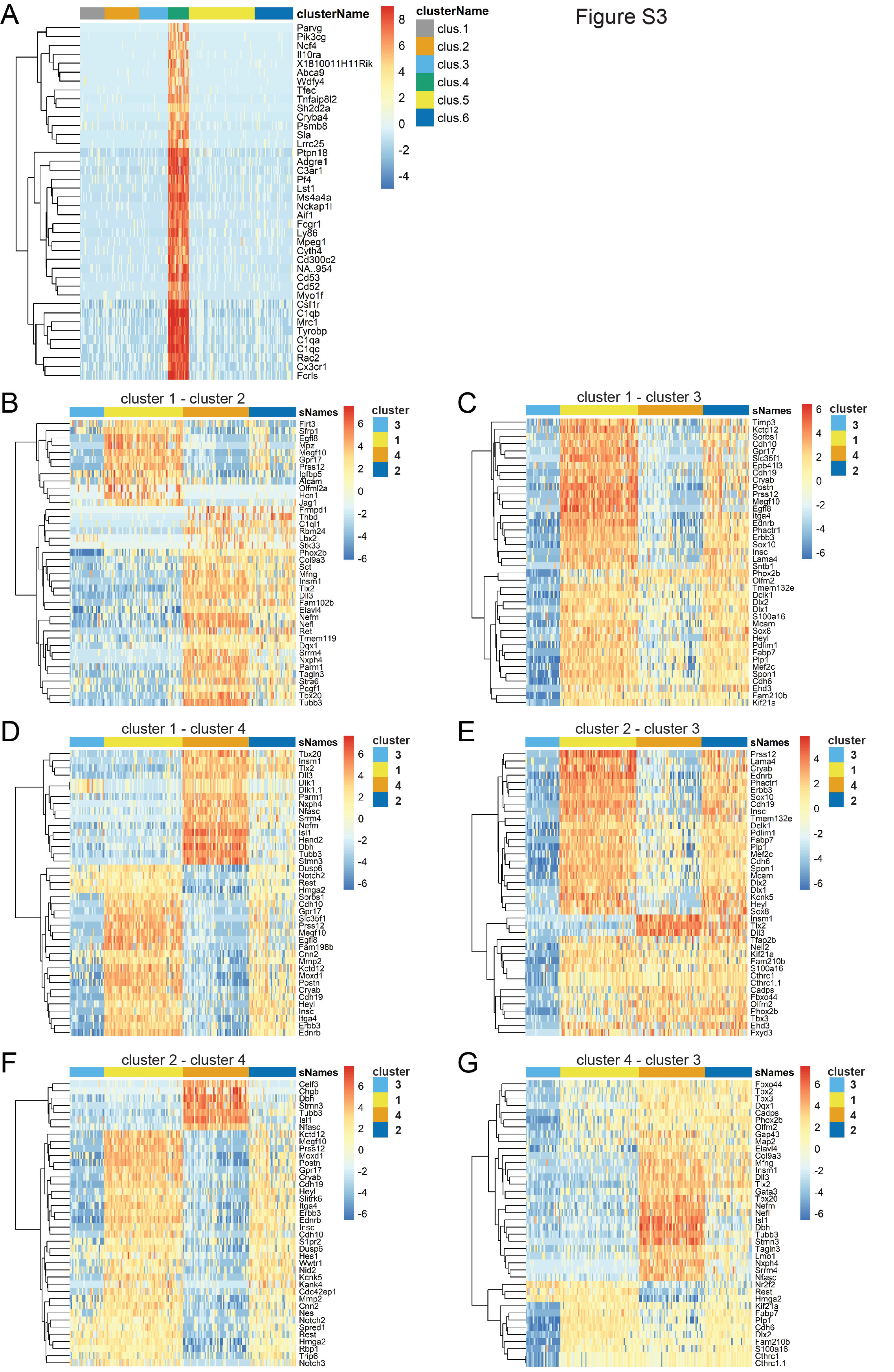

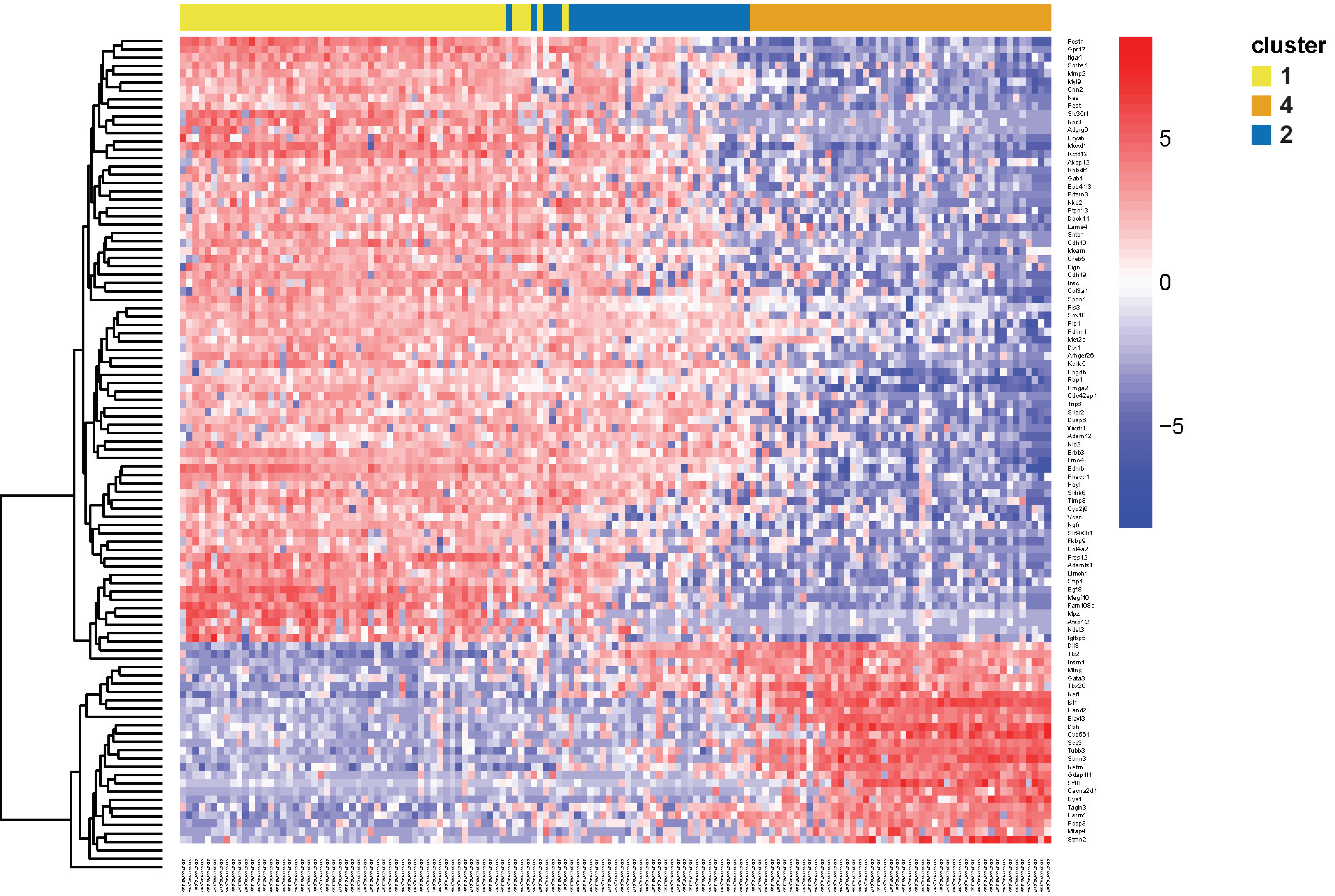


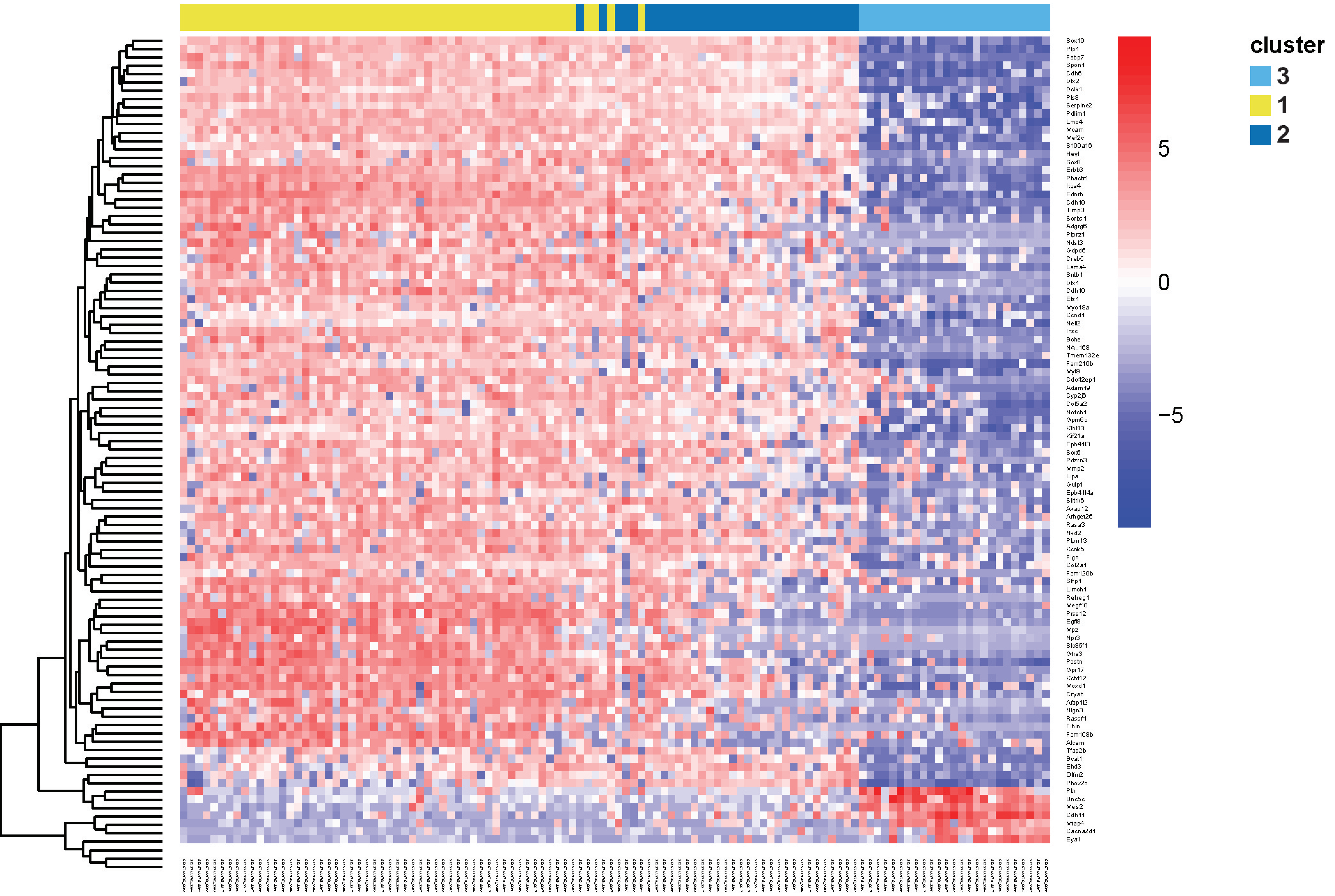


Figure S4
